# What we (don’t) know about global plant diversity

**DOI:** 10.1101/404376

**Authors:** William K. Cornwell, William D. Pearse, Rhiannon L. Dalrymple, Amy Zanne

## Abstract

**Rationale:** The era of big biodiversity data has led to rapid, exciting advances in theoretical and applied biological, ecological and conservation sciences. While large genetic, geographic and trait databases are available, these are neither complete nor random samples of the globe. Biases in species absence in these databases create problems, reducing our inferential and predictive power.

**Methods:** We performed a comprehensive examination of the taxonomic and spatial sampling in the most complete current databases for plant genes, locations, and traits.

**Results:** Only 17.7% of the world’s described land plants feature in all three databases, meaning that more than 82% of plant biodiversity lacks representation in at least one database. Species coverage is highest for location data and lowest for genetic data. Bryophytes and orchids stand out taxonomically and the equatorial region stands out spatially as poorly represented in all databases.

**Conclusion:** We have highlighted a number of clades and regions about which we know little functionally, spatially and genetically, on which we should set research targets. The scientific community should recognize and reward the significant value, both for biodiversity science and conservation, of filling in these gaps in our knowledge of the plant tree of life.

## Introduction

In perhaps the first plant trait database, Theophrastus of Eresus (371–287 BC), a student of Plato and Aristotle, assembled and published a list of the woody and herbaceous plants growing in his garden (Theophrastus, 1916). Since then the nature and scale of plant databases have changed, but the goal of organized, accessible information about the world’s plants remains the same. For the first time, after more than 2000 years of research, global versions of Theophrastus’ list are now on the horizon.

Global biological databases have two goals: (1) provide the best global snapshots of biodiversity available at a given time, and (2) organize their underlying data to facilitate their intersection with other datasets to address research questions. From decades of hard work and accumulated knowledge, we know more now than we ever have about biodiversity. The hurdles stopping us from attaining our goal of available, comprehensive global plant databases remain daunting but also increasingly surmountable. In this effort, we believe it is worthwhile, periodically, to take stock of what information we have and what we still lack. Here we ask how much do we know about global plant diversity, and what and where are the most conspicuous blank pages in our encyclopedia of plant life?

There are four primary classes of global biological data: taxonomic, geographic, genetic, and trait. Together they provide the most complete picture currently available of where and how organisms make a living across the globe and how they have evolved through time. Each of these data types is complex in its own way. Two classes, taxonomic and trait, require taxon-based specialized knowledge—botanists are required to build and curate such datasets [*e.g.*, The Angiosperm Phylogeny Website http://www.mobot.org/MOBOT/research/APweb/, The Plant List (http://www.theplantlist.org/), The State of the World’s Plants Report RBG Kew (2016), and TRY Kattge *et al.* (2011)]. In contrast, data within genetic and geographic databases share similar structure and curation protocols across all organisms and as such data for plants, animals and microbes can be usefully kept in one place. General databases for all organisms exist for genes (GenBank; Benson *et al.*, 2013), phylogenetic relationships based on those genes (The Open Tree of Life; Hinchliff *et al.*, 2015) and spatial observations (GBIF; http://gbif.org/). Here, using The Plant List as our taxonomic backbone, we explore the data available in Gen-Bank, GBIF, and TRY, as these are the most complete current databases with fundamental information about global plants.

These three databases (GenBank, GBIF, and TRY) were assembled from numerous research projects with diverse original goals and data availability; this diversity, while valid within each effort, creates global-scale biases in terms of which species feature in a particular database and which do not, which in turn effects the usability of data (Beck *et al.*, 2014, 2013; Hinchliff & Smith, 2014; Gratton *et al.*, in press). The globally uneven distribution of research funding may play a role in creating biases in data collection by limiting researchers’ efforts to certain geographical regions or taxa (particularly to species of commercial or cultural value). Furthermore, biodiversity is not evenly distributed across the globe. For instance, the high diversity, rarity, and beta-diversity of many tropical ecosystems means that the sampling effort required to capture some percentage of the species present during a sampling trip at low latitudes is much more intensive than at high latitudes. Global patterns in biodiversity, combined with patchy sampling effort, has left us with gaps in biodiversity data. Finally, non-random gaps within a given database can intersect with gaps in other databases in surprising ways, reducing the number of species about which we have a comprehensive understanding. One of our main aims is to examine these biases, gaps, and overlaps.

Such data paucity matters. Lack of data is a major impediment to conservation: we cannot save a species if we do not know where it is or what it needs to live. Data are crucial to identifying conservation crises (Cardoso *et al.*, 2011; Rodrigues *et al.*, 2006; Keith, 2015). For example, high-resolution remote-sensing deforestation maps (Hansen *et al.*, 2013), when combined with even small amounts of location data can identify rare species at risk of extinction and worthy of conservation efforts. A great deal of resources go into conservation actions for some well-studied species that are at risk; in contrast, poorly-studied species may be going extinct before conservation biologists are able to interrogate a geographic database to determine where they are found or a trait database to determine its form or characteristics should they go looking for it. Similar to recent assessments of gaps in data for birds (Team eBird, July 2014) and fungi (Nilsson *et al.*, 2016), it is our hope that an assessment of gaps in plant data across geographic space and the extant branches of the plant phylogeny will spur the sharing and/or further collection of data where they are most sorely needed.

By isolating the key spatial and taxonomic knowledge gaps in plant data, we hope to increase our chances of filling in the most important missing pages in the book of plant life as efficiently as possible. To quantify and understand these gaps we present an analysis of recent downloads from global plant databases. Here, we (1) quantify the species coverage in each of three GenBank, TRY, and GBIF databases, and (2) quantify the intersection of these databases, reporting the proportion of well-studied species (in all three databases) in contrast to patchily-studied species (in two or fewer databases) and poorly-studied species (in zero databases). Finally, we (3) identify the phylogenetic clades and geographical regions for which we have different amounts of trait and genetic data, allowing us to contrast the clades and parts of the world for which we have relatively poor and rich coverage. While plant scientists may all agree that there are gaps in our knowledge of plants, they are unlikely to know where the most gaping holes occur (FitzJohn *et al.*, 2014). By highlighting the clades and regions in most urgent need of attention, we hope to turn what appears an insurmountable task into a manageable checklist of gaps to be filled.

## Materials and Methods

Working across databases requires a common taxonomy: to combine information about species from one database with another requires a common definition of what species exist. We consider The Plant List (TPL; http://www.theplantlist.org/) to be the most complete and widely-accepted list of plant species (and their taxonomic placement) at the global scale. At the time of writing, TPL (v1.1) records 350,699 names as “accepted” species of land plants. The taxonomic scope of this analysis (and of TPL) is embryophytes (land plants), including the bryophyte grade (mosses, hornworts, and liverworts), lycophytes, monilophytes (ferns and allies), and spermatophytes (seed plants including gymnosperms and angiosperms). We emphasize that our use of TPL is conservative, as by making use only of species that are “accepted” we exclude less studied lineages in need of additional taxonomic attention. Further, variation in how sub-specific epithets are handled among databases limits our analysis to species binomials, and so we do not address cases where databases contain data on different sub-species or varieties of the same species. Continuing to resolve taxonomic controversies in a globally-accepted, versioned, and updateable list will help interoperability of these databases and benefit biodiversity science.

### Databases

We examine the state of three databases: GenBank (genetic; Benson *et al.*, 2013), the Global Biodiversity Information Facility (GBIF—geographic; http://www.gbif.org/), and TRY (functional traits; Kattge *et al.*, 2011). These databases were selected as we consider them to be the three most complete databases on plant biodiversity currently available for the most important characteristics of plants. Below we describe how each of the databases (genetic, geographic, and functional trait) were downloaded and analyzed. Each database organization does its own synonymy updating, so we did not attempt to improve on those steps.

Data were downloaded on the 21st of May, 2018. All of our code and analyses are available online (https://github.com/cornwell-lab-unsw/sTEP_overlap). The online access of our data and code allows for others to adapt our work for other purposes and permits future readers to compare how our knowledge of plants continues to change over time. We note that data quality is an issue with all three of these global databases. The quality of the data within these global collaborations continues to improve but errors remain. Improving data quality is an important and ongoing effort (Franklin *et al.*, 2017; Cicero et al., 2017; Veiga *et al.*, 2017) that will likely never end.

### Geographic data

We obtained a complete set of GBIF observations with georeferences (*i.e.*, the observation had a location associated with it) and had no “spatial issues”. Removing any records with “spatial issues” means our records were free from obvious issues (*e.g.*, latitude/longitude location swaps or mislabelled identifications over water). In our May 2018 download of GBIF we obtained 174,464,135 observations (tab-delimited file-size 101 GB). There were many incomplete identifications (e.g., *Carex sp.*) in the database, which are removed in our processing. Once these data are synchronized to the TPL species list, we are left with a total of 150,270,519 observations for 257,266 species.

### Trait data

We use the species-based summary of TRY (Kattge *et al.*, 2011) data available online (https://www.try-db.org/dnld/TryAccSpecies.txt) to obtain a list of all species for which any trait data are available in TRY. After synchronizing the species names to TPL, the database contains 124,414 species for which we have data on at least one trait.

### Genetic data

We compiled a list of all taxa listed within NCBI’s taxonomy (which is used in all GenBank searches) as green plants (Plantae). To do this, we downloaded NCBI’s complete taxonomy network, then extracted all named terminal nodes (*i.e.*, taxa) subtending from the ‘green plants’ node (number 33090 at that time). After synchronizing the species names to TPL, the database contains 108,616 species.

### Analysis

Our first and second aims are to quantify the coverage and intersection of the three databases (TRY, GenBank, GBIF). We examine the presence and absence of records for each species in TPL for each database, allowing us to take stock of the species coverage and overlap of available data. We thus identify how many species are well-studied (*i.e.* have more than zero data in all three databases), patchily-studied (*i.e.* have more than zero data in two or fewer databases) and poorly-studied (*i.e.* have zero data in all three databases). While this approach determines that species that are present in each database with at least a single record are well-studied, it does not mean that these species are particularly well represented in any given database. There is no single metric by which we could determine whether a species is ‘sufficiently’ covered in these databases, since different purposes have different data requirements; we instead focus on describing the presence or absence of any data at all.

Our third aim is to examine the taxonomic and geographic distribution patterns in sampling in all three databases. To do this, we identify the phylogenetic clades and geographical regions for which we have some trait and genetic data, allowing us to contrast the clades and parts of the world for which we have relatively poor coverage. At its core, our approach quantifies the extent of deviation from completely unbiased sampling. As with all analysis at this scale, significance testing would be of little use: all patterns described would be highly significant. Instead we seek to quantify the extent to which sampling is uneven across taxonomy and geography.

For taxonomy, we analyze all families for the degree to which sampling (species coverage) is non-randomly higher or lower than the overall rate across the entire database. To do this we apply the KSI method described in Cornwell *et al.* (2014), modified to use the G-statistic (Sokal & Rohlf, 1981) as our analysis focuses on presence/absence of data (a binary variable), unlike the continuous data in Cornwell *et al.* (2014) (code for this analysis is available at www.github.com/traitecoevo/ksi). The key feature of this approach is that it statistically balances the size of the clade with its deviation from the mean sampling rate (i.e., presence of records for species in that clade in a given database) across all land plants. Thus our results are not driven by (true) variation in the diversity of the families.

For geography, we take two approaches to determine spatial data gaps in trait and genetic databases: one considers all GBIF observations, while the other first calculates the median latitude for each species. The observation (N=150,270,519) or species (N=257,266) is then classified as either with or without trait data, with or without genetic data, and with or without both trait and genetic data. We fit six generalized additive models for very large datasets (function “BAM” within Wood, 2017) with the only predictor being latitude. The model estimates the probability of an observation or species having trait data, genetic data, or both, at a given latitude. Because fitting generalized additive models is memory intensive, even with 128 GB of RAM, random sub-sampling of the observation database down to 10,000,000 points was necessary. This sub-sampling did not affect the result, however, with repeated samples producing identical fits.

To discover proportionally under-sampled families by biogeographic region, we split the raw GBIF observations by continent (Environmental Research Systems Institute, ESRI). We isolate species that are only found in one biogeographic region, that is, species endemic to that particular region. Considering each region separately, we use the KSI approach to quantify the most disproportionately under-sampled families within each region. This approach was designed to identify region-specific gaps in data. Note that this analysis excludes both species with distributions on multiple continents and species that are invasive somewhere, but we expect these species to be in the well-studied category.

## Results

Of the accepted plant species of the world, 17.7% (61,902 species) are well-studied, meaning we know *something* about their genetic make-up, *something* about their functional traits, and *something* about their geo-referenced location (Table 1). These species make up the “well-studied” section of the Venn diagram (Figure 1), where genetic data, trait data and geographic data overlap.

**Table 1:** Count of accepted plant species in each of the database and overlap of species among two- and three-way combinations of databases.

|  | Genbank | GBIF | TRY | All three |
| --- | --- | --- | --- | --- |
| Genbank | 108616 | 100507 | 63783 |  |
| GBIF | 100507 | 257266 | 114067 |  |
| TRY | 63783 | 114067 | 124414 |  |
| All three |  |  |  | 61902 |

**Figure 1:**
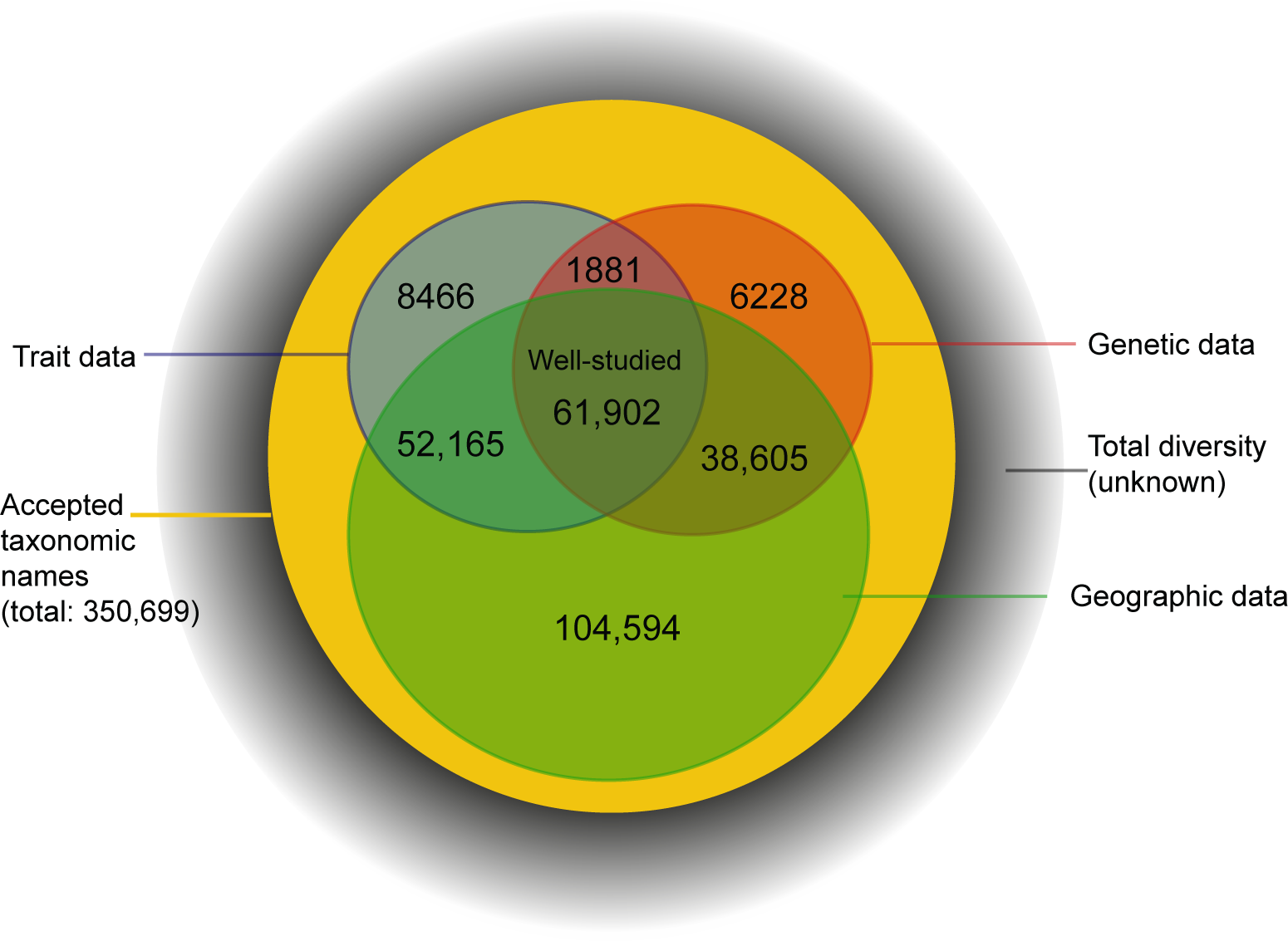
Shortfalls in our knowledge of the plant kingdom. (1)The Linnean shortfall (diversity for which we have no names) is the area outside of the “accepted taxonomic names” footprint. (2)The Darwinian shortfall (diversity for which we have zero genetic information in GenBank) is the area outside of the “Genetic data” footprint. (3)The Wallacean shortfall (diversity for which we have zero location information in GBIF) is the area outside of the “Geographic data” footprint.(4)The Warming shortfall (diversity for which we have zero records of functional trait data in TRY) is the area outside of our “Trait data” coverage. (5)The Venn shortfall, a culmination of these first four shortfalls, is the area outside of where these four kinds of knowledge intersect - the area over overlap where species are considered “well-studied”.

More than half of all plant species - 55.6% - are patchily-studied across the datasets, meaning they are present in two or fewer datasets (Table 1; Figure 1). Building on centuries of collections and a global network of museums and herbaria, GBIF has by far the most complete species coverage, with occurrence data for 257,266 species (73.3% of accepted diversity). In TRY, trait data are available for 124,414 species (35.5% of accepted diversity). The database with the least coverage is GenBank, where genetic data are available for 108,616 species (31.0% of accepted diversity).

For 26.7% of plant species, we know nothing of them but their names. These species are poorly-studied and we have no genetic, location or functional trait data in the databases considered (Figure 1).

### Where we know a lot

All large data sources have areas of strength and weakness across the plant phylogeny. There are several hotspots in our knowledge, and the families that are well-studied (that is, having higher than average proportion of member species occupying the “well-studied” section of Figure 1) tend to be widespread and of economic importance either as food or resource crops, or as cultivars (Table 3; Figure 3). These include the Poaceae (grasses, bamboos), Pinaceae (cedars, pines, spruces), Solanaceae (nightshades, with tomatoes, potatoes, peppers and eggplants), Moraceae (mulberries, breadfruit, jackfruit, figs), Eurphorbiaceae (rubber, spurge cultivars), Proteaceae (Macadamia, silky oak, cultivars), Iridaceae (irises, saffron) and Polemonaceae (Phlox). There are well-studied families which do not have obvious commercial value but may simply have been of interest for research and exploration by particular research groups: two that came out in the top 10 are the Zamiaceae (cycads) and Orobanchaceae (broomrapes, mostly parasitic). Other areas of the phylogeny with good sampling across all datasets are the gymnosperms and early diverging angiosperms (Figure 3), perhaps because they were identified by Darwin as informative with respect to the “abominable mystery” of the origin of angiosperms. Some of the orders with high coverage (Figure 3) are that way because of their unusually small clade size. For instance, Trochodendrales and Acorales have full coverage — both orders are comprised of only two accepted species, while Ginkgoales consists of a single accepted species.

**Table 2:** The three most undersampled families for species endemic to each continent. Note that endemicity in this analysis is based on gbif observations, and may be a underestimate due to species that are invasive elsewhere.

| Continent | Database | Family | Proportion of species sampled | Estimated continental endemics | Distinctiveness index |
| --- | --- | --- | --- | --- | --- |
| Africa | TRY | Acanthaceae | 0.097 | 963 | 227.263 |
|  | TRY | Asteraceae | 0.195 | 2513 | 137.427 |
|  | TRY | Aizoaceae | 0.149 | 955 | 113.331 |
|  | Genbank | Acanthaceae | 0.173 | 963 | 87.737 |
|  | Genbank | Begoniaceae | 0.009 | 111 | 70.565 |
|  | Genbank | Lauraceae | 0.061 | 163 | 59.395 |
| Asia | TRY | Orchidaceae | 0.050 | 3085 | 1094.982 |
|  | TRY | Gesneriaceae | 0.028 | 617 | 269.321 |
|  | TRY | Asteraceae | 0.150 | 2486 | 216.418 |
|  | Genbank | Orchidaceae | 0.284 | 3085 | 149.851 |
|  | Genbank | Begoniaceae | 0.051 | 234 | 144.026 |
|  | Genbank | Rubiaceae | 0.247 | 1352 | 118.264 |
| Australia | TRY | Orchidaceae | 0.079 | 557 | 295.071 |
|  | TRY | Lamiaceae | 0.149 | 261 | 72.413 |
|  | TRY | Araceae | 0.024 | 42 | 32.383 |
|  | Genbank | Orchidaceae | 0.217 | 557 | 56.275 |
|  | Genbank | Frankeniaceae | 0.000 | 32 | 28.407 |
|  | Genbank | Rubiaceae | 0.144 | 111 | 25.815 |
| Europe | TRY | Asteraceae | 0.262 | 1382 | 92.533 |
|  | TRY | Orchidaceae | 0.103 | 213 | 79.508 |
|  | TRY | Lamiaceae | 0.192 | 193 | 28.876 |
|  | Genbank | Asteraceae | 0.158 | 1382 | 258.282 |
|  | Genbank | Rosaceae | 0.197 | 615 | 54.249 |
|  | Genbank | Onagraceae | 0.000 | 24 | 18.678 |
| North America | TRY | Crassulaceae | 0.181 | 215 | 119.096 |
|  | TRY | Pottiaceae | 0.000 | 47 | 73.190 |
|  | TRY | Bryaceae | 0.000 | 46 | 71.631 |
|  | Genbank | Myrtaceae | 0.064 | 297 | 154.550 |
|  | Genbank | Begoniaceae | 0.009 | 109 | 90.513 |
|  | Genbank | Primulaceae | 0.093 | 215 | 87.264 |
| NZ and Oceania | TRY | Orchidaceae | 0.077 | 195 | 84.651 |
|  | TRY | Rubiaceae | 0.153 | 235 | 48.578 |
|  | TRY | Pandanaceae | 0.051 | 59 | 30.951 |
|  | Genbank | Phyllanthaceae | 0.098 | 82 | 29.027 |
|  | Genbank | Euphorbiaceae | 0.059 | 51 | 25.326 |
|  | Genbank | Orchidaceae | 0.200 | 195 | 22.924 |
| South America | TRY | Orthotrichaceae | 0.000 | 128 | 182.369 |
|  | TRY | Pottiaceae | 0.000 | 127 | 180.941 |
|  | TRY | Dicranaceae | 0.000 | 109 | 155.250 |
|  | Genbank | Rubiaceae | 0.148 | 1923 | 174.428 |
|  | Genbank | Begoniaceae | 0.018 | 219 | 107.190 |
|  | Genbank | Araceae | 0.129 | 848 | 103.845 |

**Table 3:** The most well-sampled families for each database, as well as those which are well-sampled across all three databases, i.e., having a disproportionately high proportion of species in the “overlap” section of Figure 1.

| Database | Family | Proportion<br>sampled | Estimated<br>diversity | Distinctiveness<br>index |
| --- | --- | --- | --- | --- |
| TRY | Poaceae | 0.77 | 11470 | 8648.92 |
|  | Begoniaceae | 0.96 | 1601 | 2754.26 |
|  | Euphorbiaceae | 0.67 | 6547 | 2670.13 |
|  | Orobanchaceae | 0.91 | 1614 | 2218.14 |
|  | Santalaceae | 0.90 | 992 | 1302.79 |
|  | Moraceae | 0.85 | 1217 | 1263.58 |
|  | Loranthaceae | 0.91 | 887 | 1185.14 |
|  | Urticaceae | 0.74 | 1465 | 895.42 |
|  | Proteaceae | 0.76 | 1253 | 854.95 |
|  | Iridaceae | 0.61 | 2315 | 644.84 |
| Genbank | Brassicaceae | 0.53 | 4065 | 895.56 |
|  | Restionaceae | 0.84 | 529 | 635.86 |
|  | Apiaceae | 0.52 | 3270 | 631.14 |
|  | Liliaceae | 0.65 | 745 | 365.97 |
|  | Pinaceae | 0.87 | 255 | 349.81 |
|  | Poaceae | 0.39 | 11470 | 339.32 |
|  | Zamiaceae | 0.88 | 216 | 312.67 |
|  | Polemoniaceae | 0.71 | 455 | 302.12 |
|  | Plantaginaceae | 0.51 | 1614 | 296.57 |
|  | Ericaceae | 0.44 | 3553 | 282.12 |
| GBIF | Proteaceae | 0.96 | 1253 | 458.72 |
|  | Solanaceae | 0.88 | 2677 | 337.89 |
|  | Myrtaceae | 0.83 | 5970 | 335.46 |
|  | Ericaceae | 0.85 | 3553 | 258.46 |
|  | Passifloraceae | 0.93 | 932 | 247.73 |
|  | Malvaceae | 0.83 | 4506 | 232.37 |
|  | Chrysobalanaceae | 0.97 | 535 | 230.45 |
|  | Melastomataceae | 0.83 | 4078 | 226.11 |
|  | Iridaceae | 0.86 | 2315 | 222.26 |
|  | Piperaceae | 0.85 | 2658 | 217.26 |
| All three | Poaceae | 0.31 | 11470 | 1319.29 |
|  | Pinaceae | 0.84 | 255 | 529.78 |
|  | Euphorbiaceae | 0.27 | 6547 | 385.33 |
|  | Orobanchaceae | 0.38 | 1614 | 382.65 |
|  | Solanaceae | 0.33 | 2677 | 373.78 |
|  | Moraceae | 0.41 | 1217 | 365.32 |
|  | Polemoniaceae | 0.56 | 455 | 333.91 |
|  | Zamiaceae | 0.71 | 216 | 296.21 |
|  | Iridaceae | 0.32 | 2315 | 287.26 |
|  | Proteaceae | 0.37 | 1253 | 274.77 |

The spatial patterns in genetic and trait data availability (Figure 2) are evidence of the non-random distribution of research sites and unequal sampling efforts across latitudes. There are more data available in GenBank and TRY for the northern hemisphere than the southern hemisphere and both temperate regions are better sampled than the equatorial region (Figure 2). The availability of both trait and genetic data is best around 45 ° N. If you were to take a random GBIF observation from that latitude you would have a greater than 95% chance of accessing trait data for that species, a 90% chance of there being genetic data, and almost 90% chance of there being both genetic and trait data available. If you were to select a species at random at this latitude, there would be a 65% chance of there being trait data, a 60% chance of there being genetic records for this species, but only 45% chance of both data types being available (Figure 2).

**Figure 2:**
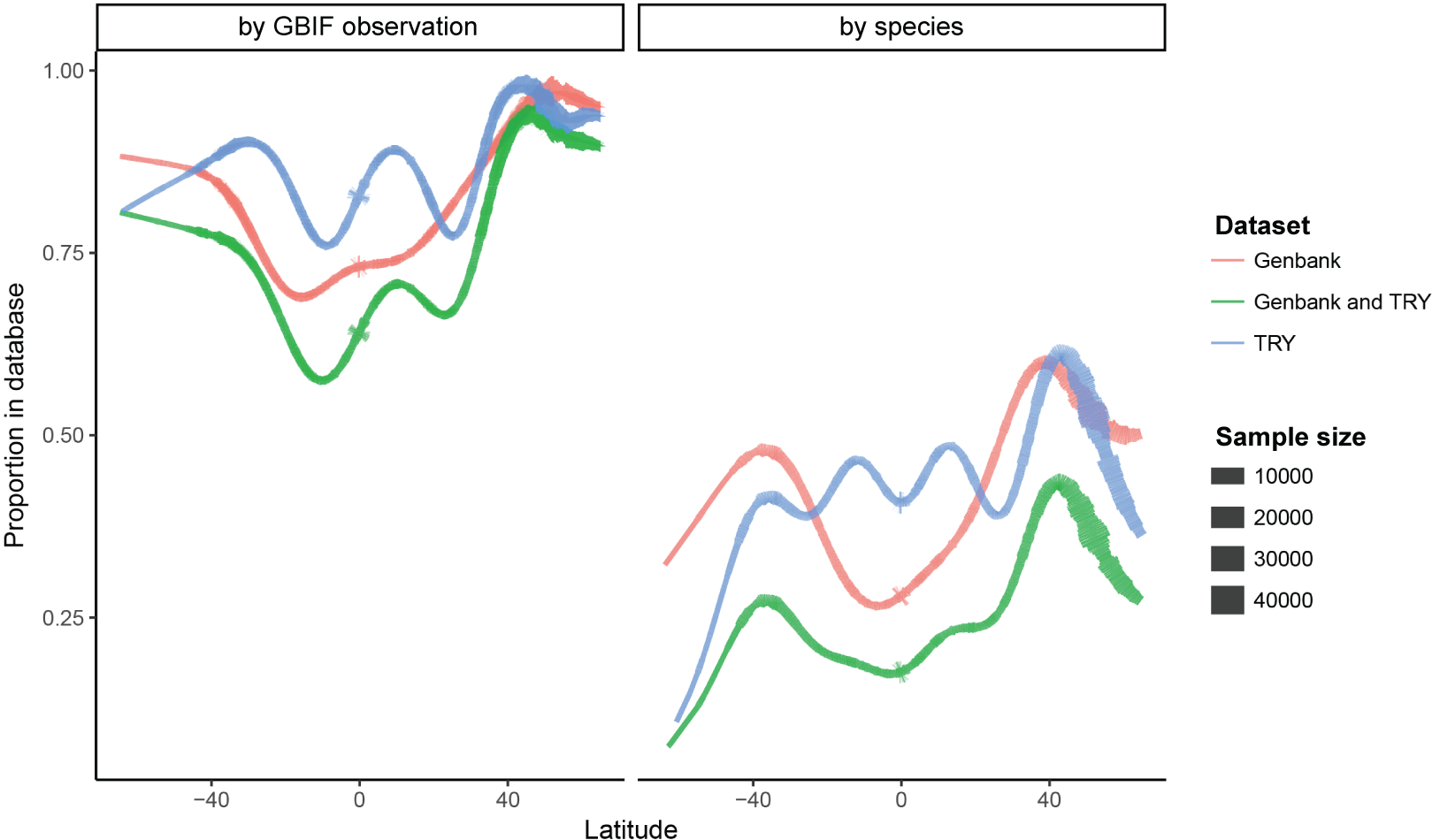
Generalized additive models for the relationship between latitude and availability of plant data. The left panel models individual GBIF observations (N=150, 270, 519); the right panel models species median values for latitude (N=257, 266). Line thickness is indicative of sample size. Note that tropical areas (esp. just south of the equator) have the poorest data coverage, and just north of 40° N is where we find the best coverage of both functional trait and genetic plant data.

**Figure 3:**
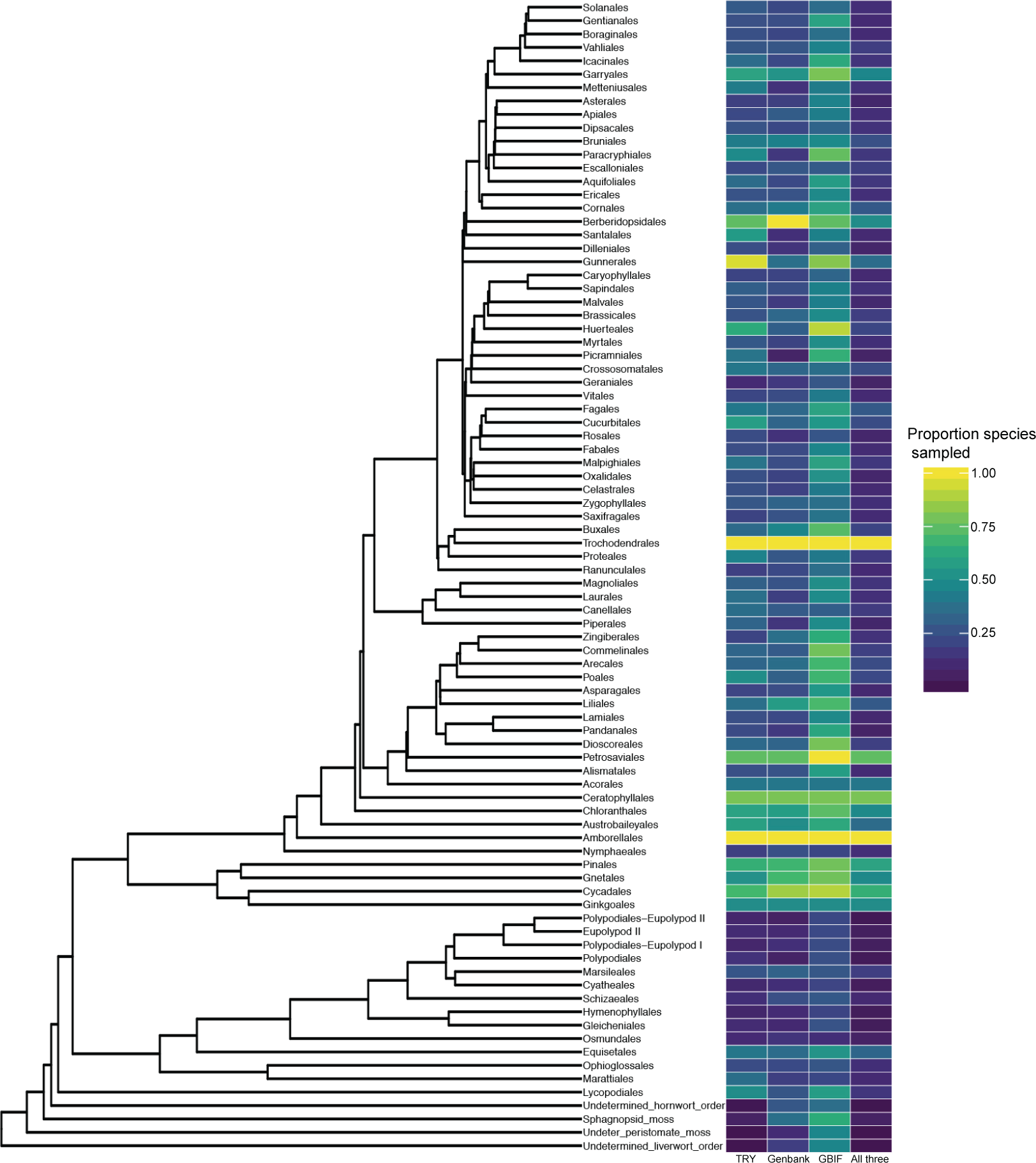
The proportion of species in each order of the tree of life for which we have functional trait, genetic and location data, and the proportion which are well-sampled, having all three types of data available.

For many accepted species, there are 10 or fewer geographical observance records and no genetic or trait data (Figure 4). The species for which there are upwards of 1000 spatial records in GBIF include the dominant trees and shrubs of North America, Europe, and South East Australia. As these species are prevalent in regions where many scientists live and work, they are well documented in numerous way and make up much of the “well-studied” section of the Figure 1.

**Figure 4:**
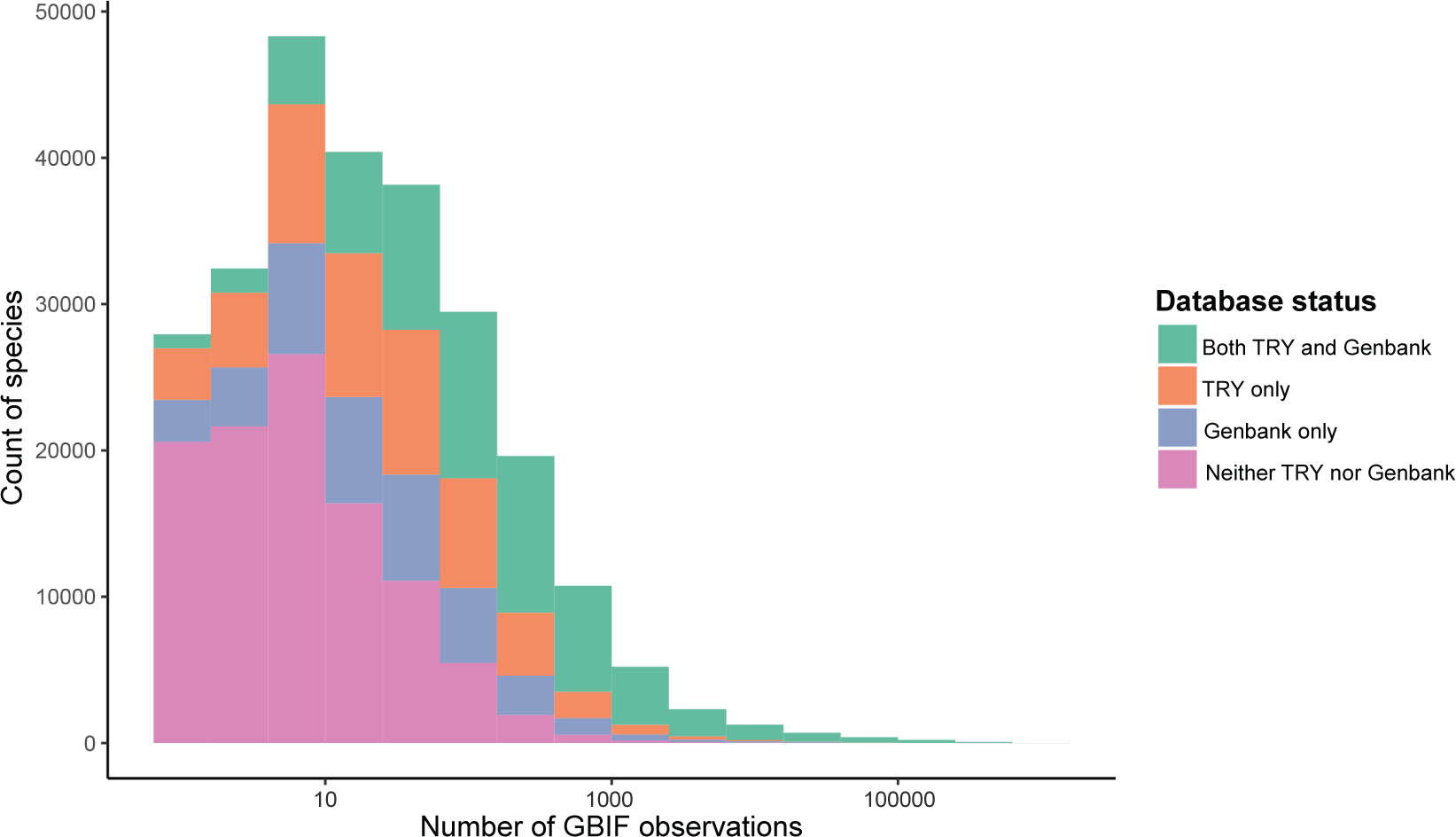
Ranking of the number of occurrences in GBIF for plant species; colors indicate the presence of trait and genetic data. Species with many spatial records also tend to have genetic and trait data available. Species with few GBIF records are less likely to be in TRY or GenBank. Note the log-scaled x-axis.

### Where we don’t know a lot

While the databases all lack a certain number of species contributing to the known extant diversity, there are plant families for which we have little data across all databases (Table 4, Figure 3). Most of the poorly-studied families, those families that are made up of species that are undersampled geographically, genetically, *and* functionally, are moss or liverwort families, but Orchidaceae, Asteraceae and Begoniaceae also earn this title. When we consider the taxonomic weaknesses of databases individually, we find that they shar similarities in the distribution of data gaps (Table 4). GBIF is extremely poorly sampled for mosses and liverworts—all of the top ten most geographically undersampled families come from these two clades. Moss and liverwort families are also undersampled for both traits and genes, nine and seven of the top ten most undersampled families in these data sets come from these clades, respectively. The traits of Orchidaceae remain extremely undersampled despite its widespread cultivation and intense horticultural use. For genetic data, the other families that make the top ten most undersampled list are all fairly diverse and widespread tropical understory families: Begoniaceae, Acanthaceae, and Piperaceae.

**Table 4:** The most disproportionately undersampled families within each database. Those families which are undersampled across the board are the “least-scienced”, having the lowest proportion of Representatives in the “overlap” section the Venn diagram.

| Database | Family | Proportion<br>sampled | Estimated<br>diversity | Distinctiveness<br>index |
| --- | --- | --- | --- | --- |
| try | Orchidaceae | 0.19 | 27782 | 4131.81 |
|  | Pottiaceae | 0.0003 | 3169 | 2775.82 |
|  | Hypnaceae | 0.0004 | 2519 | 2200.85 |
|  | Lejeuneaceae | 0.0004 | 2269 | 1980.10 |
|  | Bryaceae | 0 | 2103 | 1849.77 |
|  | Dicranaceae | 0.003 | 2116 | 1775.56 |
|  | Sematophyllaceae | 0.001 | 1629 | 1403.18 |
|  | Orthotrichaceae | 0.002 | 1272 | 1078.43 |
|  | Brachytheciaceae | 0.005 | 1102 | 909.56 |
|  | Hookeriaceae | 0.002 | 831 | 703.53 |
| genbank | Orchidaceae | 0.22 | 27782 | 1179.95 |
|  | Hypnaceae | 0.05 | 2519 | 1048.10 |
|  | Pottiaceae | 0.09 | 3169 | 902.62 |
|  | Begoniaceae | 0.02 | 1601 | 899.25 |
|  | Bryaceae | 0.07 | 2103 | 753.64 |
|  | Sematophyllaceae | 0.09 | 1629 | 457.60 |
|  | Dicranaceae | 0.12 | 2116 | 423.91 |
|  | Orthotrichaceae | 0.10 | 1272 | 316.40 |
|  | Fissidentaceae | 0.07 | 817 | 289.72 |
|  | Acanthaceae | 0.19 | 3948 | 277.58 |
| gbif | Orchidaceae | 0.57 | 27782 | 3833.24 |
|  | Hypnaceae | 0.40 | 2519 | 1249.90 |
|  | Pottiaceae | 0.46 | 3169 | 1058.46 |
|  | Sematophyllaceae | 0.42 | 1629 | 703.30 |
|  | Bryaceae | 0.46 | 2103 | 695.73 |
|  | Dicranaceae | 0.52 | 2116 | 455.56 |
|  | Lejeuneaceae | 0.54 | 2269 | 394.40 |
|  | Brachytheciaceae | 0.49 | 1102 | 288.83 |
|  | Neckeraceae | 0.46 | 816 | 273.87 |
|  | Fissidentaceae | 0.49 | 817 | 217.23 |
| All three | Orchidaceae | 0.09 | 27782 | 1848.76 |
|  | Pottiaceae | 0.0003 | 3169 | 1214.53 |
|  | Hypnaceae | 0.0004 | 2519 | 961.78 |
|  | Lejeuneaceae | 0.0004 | 2269 | 864.75 |
|  | Bryaceae | 0 | 2103 | 814.59 |
|  | Dicranaceae | 0.002 | 2116 | 764.57 |
|  | Sematophyllaceae | 0.0006 | 1629 | 616.81 |
|  | Orthotrichaceae | 0.0008 | 1272 | 478.85 |
|  | Begoniaceae | 0.02 | 1601 | 423.88 |
|  | Asteraceae | 0.14 | 32920 | 405.18 |

We identify the three family groups for which we have the greatest gaps in data for endemic species of each of the different continents (Table 2). As well as seeing similar overall patterns in data gaps to those outlined above, there are some families which made multiple appearances. More data are needed for Rubiaceae endemics of Asia, Australia, NZ and Oceania, and South America; for Asteraceae endemics of Europe, Asia and Africa; for Lamiaceae endemics of Europe and Australia; and for Araceae endemics of Australia and South America.

The low latitudes are hotspots for plant diversity, home to some of the most diverse ecosystems on the planet, and yet we find ourselves with a paucity of plant data for these tropical regions. It is in the equatorial region where we find the least species coverage of genetic and trait data. For instance, if one were to select a single species at random at a latitude of 0°, there would only be a 40% chance of data for that species being available in TRY, a mere 27% chance of there being data for that species in GenBank (dotted lines; Figure 2), and less than a 20% chance of both trait and genetic data being available.

## Discussion

### Shortfalls in our knowledge of global plant diversity

Our knowledge of plants is incomplete in a number of well-defined ways (see Hortal *et al.*, 2015), four of which are pertinent to the database resources we have analyzed here. First, given the rate at which species are being discovered, we are quite confident that we have not sampled and described all plant life on Earth—this is the *Linnean shortfall* in our knowledge of biodiversity (Lomolino, 2004). Second, we lack basic taxonomic or phylogenetic information on the placement of many plants in the tree of life—this is the *Darwinian shortfall* (Diniz-Filho *et al.*, 2013). Third, we do not know the distribution (range) of many plants on Earth—this is the *Wallacean shortfall* (Lomolino, 2004). Fourth, we do not know about the functional traits of many plants—this is the *Raunkiæran shortfall* (Hortal *et al.*, 2015). A major result from our study is that the Wallacean shortfall (26.7%) is currently far less severe than the Darwinian (69%) or Raunkiæran (64.5%) shortfalls.

Methods within some lines of scientific inquiry lend themselves well to systematic sampling. Remote sensing, for example, surveys every pixel of the planet for an extensive range of abiotic and biotic features. Similarly, despite the uneven distribution of biodiversity and the challenge in getting to remote areas, taxonomy has an inherent goal to describe every species on earth. That the Wallacean shortfall is less severe than others is due mainly to its strong link to taxonomy related resources — GBIF has had great success in encouraging herbaria and museum collections to participate in systematic centralization of data from collections; at the time of writing, a 57th country had joined the initiative. Nonetheless, there is more to do: the gap in our geographic records of the plant kingdom currently stands at 26.7%, meaning that there are 93,433 accepted species for which we do not know even a single location in the world they grow or have grown. This is literally lost diversity data—for the species to have been described taxonomically, it was collected, but the geographic information (the location where that specimen was collected) been lost.

As we lack important information about where they live, we also then lack information about their ecology and evolution. Most worryingly, without even basic geographic information, any true assessment of the conservation status of these species is impossible. However, as more of the world’s herbarium collections become digitized and uploaded directly to GBIF, or through intermediary initiatives, we do expect that this shortfall will decline relatively rapidly. Indeed, comparison of the current coverage of Gen-Bank, GBIF and TRY to that from a preliminary analysis for this paper in 2015 indicated that the Wallacean shortfall is decreasing the fastest (see Appendix 1).

Relative to the Wallacean shortfall, the Raunkiæran and Darwininan shortfalls are more severe. The explicit goal of complete sampling of plant diversity for either trait or genetic data has never been clearly stated perhaps because it was so distant, and the result is patchy species coverage. The Darwinian shortfall is the most severe of the three: we have a complete lack of genetic information for 69% of known plant species. Worryingly, this dataset has shown the slowest pace of growth in species coverage over the last 3 years (see Appendix 1), likely because it is much more laborious and expensive to produce a single record of genetic information than for either of the other databases. The Raunkiæran shortfall stands at 64.5% of accepted diversity, which is a considerable gap in our plant knowledge given the current focus on trait-based analyses in the scientific exploration of ecological and evolutionary processes. There are other taxonomic groups, including birds and mammals, for which complete sampling has been an explicit aim, and as such relevant trait datasets have high-to-complete species coverage (e.g. Wilman *et al.*, 2014; Jones *et al.*, 2009). Despite the daunting diversity of land plants, we are now at the point where complete sampling through targeted campaigns is a feasible goal.

Perhaps the most worrying of these shortfalls is the Linnean as it is hard to know anything about what we do not know; these are the species for which we know literally nothing and they do not even have a name. Estimations are that the number of un-named plant species may be 10-20% of the number of known plant species (Joppa *et al.*, 2011) and that the majority may inhabit diverse, tropical regions (Wheeler *et al.*, 2004). These may have never been collected (collection of new species in the field may be the greatest rate-limiting step towards mapping the whole tree of plant life (May, 2004)), or they may have been collected but mis-identified or are yet to be described. If they have been collected, there may be associated genetic, functional and spatial data but without a name as a link, this information is lost to us. It can be difficult for taxonomists to keep up with the influx of new specimens needing description (and the revisions that new specimens can warrant), and there may not be the expertise or financial support available to do so-often resulting in significant lag time between collection and publication of descriptions with accepted scientific names (Bebber *et al.*, 2010; Wheeler *et al.*, 2004). With new information and taxonomic effort the number of accepted species will no doubt rise in the coming years (see Mora *et al.*, 2011); however, the training of taxonomists and the financial support of their research in both new and existing collections is crucial to the reduction of the Linnean shortfall.

Each of these four gaps in our knowledge of plant diversity combines to create what we suggest is a fifth shortfall in our knowledge of global plant diversity: we do not know *anything* about the geography, *anything* about the genes, nor *anything* about the traits for the vast majority of known plant species. We term this the *Venn shortfall*, since it relates to a lack of intersection among databases. The Venn shortfall is a critical issue that affects both biodiversity science and conservation. How can we assess conservation risk of a species if all the information we have about them is a name? The narrow overlap among the different dimensions of our knowledge of plant diversity means that much research and policy-decision making is limited to data on a relatively small (17.7% of accepted diversity) and non-random subset of plants species, and we know that when data are biased or extremely sparse we are at risk of missing or misidentifying patterns or processes(Hortal *et al.*, 2015). If we acknowledge the Venn shortfall as a serious problem facing conservation biodiversity science, then we can treat this as an important goal worthy of attention and funding.

### Closing the gaps

While there can never be a complete substitute for real data, temporary solutions have been proposed to leverage existing data to circumvent the shortfalls we describe above. These include *phyndr*, an algorithm that maximizes the overlap between phylogenetic and species’ data (i.e., trait and spatial Pennell *et al.*, 2016), *Phylomatic* which uses taxonomic data to fill-in missing phylogenetic data, and approaches which use taxonomic information to “gap-fill” our knowledge of plant functional diversity (Schrodt *et al.*, 2015; FitzJohn *et al.*, 2014; Taugourdeau *et al.*, 2014). As human pressure on plant diversity increases through time, such techniques may be our best hope of estimating policy-relevant information immediately but they also introduce uncertainty, and dealing with this uncertainty in downstream analyses may be non-trivial. Leaving aside the difficulties involved in imputing data and then going out and collecting it, most simple models of trait evolution for plants are far from adequate (Pennell *et al.*, 2015) and heterogeneity in evolutionary processes among the branches of the tree is common (Eastman *et al.*, 2011)). Such evolutionary methods can be powerful for filling in small gaps where information on a species can be estimated based on its close relatives, but are thus of limited use in large clades with poor data coverage. Most worryingly, because gap-filling in most cases uses taxonomic or phylogenetic information to infer species values, there is the danger of circularity in many contexts, such as evolutionary and biogeographic studies. For goals that are specific to the unsampled species—such as, what is the species’ ecology and its conservation risk—imputation will simply not be informative given the current patchy data. Such approaches cannot supersede the collection of novel data, but they do help us make the most of what data we do have.

Unlike the Linnean shortfall, targeted programs may reduce the magnitude of the Venn, Darwinian, Wallacean and Raunkiæran shortfalls. Held within the world’s > 3000 herbaria are almost 390 million specimens(Thiers, 2017) and considerable efforts are underway to increase availability of data from these collections through large digitization and sharing initiatives (Pyke & Ehrlich, 2010; James *et al.*, 2018). The number of species for which we have names but no other data is large (13.2%), however, these species *must* have had museum/herbarium records at some time. If they still exist and are findable, these specimens contain sufficient data to address multiple shortfalls at once: there are collection location data (albeit for older specimens this tends to be at a course scale (Pyke & Ehrlich, 2010)), and the vouchers themselves can be sources of genetic and trait data (Younis *et al.*, 2018; Corney *et al.*, 2012; Li *et al.*, 2015). To some extent this is happening, but current progress is haphazard, and to be maximally efficient with our sampling efforts, it is worth (1) prioritizing herbaria that are known to have collections of undersampled taxa, especially those that are missing from all three of the databases, and (2) gathering data from specimens in locations known to be effected by habitat loss as quickly as possible (as these specimens might be the only collections that we ever obtain).

Our survey of global plant diversity leads us to make four observations that can inform a more optimal strategy for filling in knowledge gaps. First, grant programs can have direct impacts on the future of the shortfalls we have demonstrated by incentivizing work which creates overlapping species data, and such opportunities should be expanded. We suggest that funding programs should support researchers collecting and sharing data on undersampled regions or clades of the world. Second, publication of results from genetic data has long required centralized deposition of raw sequences with meta-data; A similar approach within collectors of locations and trait data would broaden the base of contributors and rapidly increase availability of both spatial and trait data. Third, one clear bias is that many well-studied families are those with some commercial value. However, species do not need to have direct commodity or cultural usefulnesses to be important (Ehrlich & Wilson, 1991). There is immense basic and applied value in novel data on undersampled families (orchids, Begoniaceae, Asteraceae, and most bryophyte families see Table 4) in addressing biases in our datasets to future biodiversity and conservation science. Last, we need to intensify genetic and trait sampling efforts in undersampled regions: spatial biases are evident in the availability of both genetic and functional trait data.

### Conclusions

Despite shortfalls (Linnaen, Darwinian, Wallacean, Raunkiæran, Venn) within and across data sources, there are more reasons than ever to be optimistic about the future of our knowledge of global plant diversity. Biodiversity sciences are progressing in leaps and bounds, empowered by the sharing of large amounts of data. However, the intersection of data among our plant databases is particularly patchy and non-random, and as we move towards a more complete picture of global functional diversity, more data are clearly needed. By taking a targeted approach to expanding and improving the status of our data foundations for such work we can progress towards an ever more complete picture of the many facets of plant biodiversity, and greater inferential and predictive power from our global plant data.

Gap-filling is also a goal among those who study other taxa: interestingly eBird — an ornithologic citizen science effort — performed a gap analysis similar to ours in 2014 and were able to generate gap-filling enthusiasm, which led to the sharing of data on 227 out of the 421 missing bird species (see https://ebird.org/news/ebirds-missing-species/ and https://ebird.org/news/ebirds-missing-species-v2/). The shear number of missing species is much larger for plants, but publicizing target lists is a good first step, and with enough of a reward, progress could be rapid. Let us begin!

## Acknowledgment

This research includes computations using the Linux computational cluster Katana supported by the Faculty of Science, UNSW Sydney. This research was initiated as part of the sTEP working group funded by sDIV, the Synthesis Centre of the German Centre for Integrative Biodiversity Research Halle-Jena-Leipzig.

## Author contributions

WKC and AEZ conceived of the initial ideas and questions. WKC and WP designed and performed the analyses. All authors wrote and revised the manuscript.

## Appendix 1

Preliminary work for this manuscript began in March of 2015. At that time data were downloaded in a similar manner to that presented in the main document. Table S1 shows the species coverage of each dataset at that time. While the data downloads of 2015 and 2018 are not perfectly comparable (due to small changes in data handling/cleaning both by us and by the databases), the changes in species coverage over time are indicative of the relative rate at which shortfalls in our knowledge of plant diversity are being addressed.

**Table S1:**
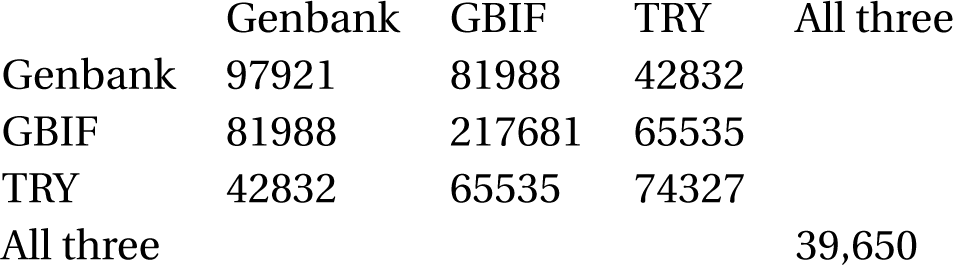
Coverage of accepted plant species in the three databases in data downloads from March 2015

|  | Genbank | GBIF | TRY | All three |
| --- | --- | --- | --- | --- |
| Genbank | 97921 | 81988 | 42832 |  |
| GBIF | 81988 | 217681 | 65535 |  |
| TRY | 42832 | 65535 | 74327 |  |
| All three |  |  |  | 39,650 |

## References

Bebber DP, Carine MA, Wood JRI, Wortley AH, Harris DJ, Prance GT, Davidse G, Paige J, Pennington TD, Robson NKB et al. 2010. Herbaria are a major frontier for species discovery. Proceedings of the National Academy of Sciences, 107: 22169–22171.

Beck J, Ballesteros-Mejia L, Nagel P, Kitching IJ. 2013. Online solutions and the ‘wallacean shortfall’: what does gbif contribute to our knowledge of species’ ranges? Diversity and Distributions, 19: 1043–1050.

Beck J, Böller M, Erhardt A, Schwanghart W. 2014. Spatial bias in the gbif database and its effect on modeling species’ geographic distributions. Ecological Informatics, 19: 10–15.

Benson DA, Cavanaugh M, Clark K, Karsch-Mizrachi I, Lipman DJ, Ostell J, Sayers EW. 2013. Genbank. Nucleic Acids Research, 41: D36–D42.

Cardoso P, Erwin TL, Borges PA, New TR. 2011. The seven impediments in invertebrate conservation and how to overcome them. Biological Conservation, 144: 2647–2655.

Cicero C, Spencer CL, Bloom DA, Guralnick RP, Koo MS, Otegui J, Russell LA, Wieczorek JR. 2017. Biodiversity informatics and data quality on a global scale1. The Extended Specimen: Emerging Frontiers in Collections-Based Ornithological Research.

Corney DP, Clark JY, Tang HL, Wilkin P. 2012. Automatic extraction of leaf characters from herbarium specimens. Taxon, 61: 231–244.

Cornwell WK, Westoby M, Falster DS, FitzJohn RG, O’Meara BC, Pennell MW, McGlinn DJ PB et al. 2014. Functional distinctiveness of major plant lineages. Journal of Ecology, 102: 345–356.

Diniz-Filho JAF, Loyola RD, Raia P, Mooers AO, Bini LM. 2013. Darwinian shortfalls in biodiversity conservation. Trends in Ecology & Evolution, 28: 689–695.

Eastman JM, Alfaro ME, Joyce P, Hipp AL, Harmon LJ. 2011. A novel comparative method for identifying shifts in the rate of character evolution on trees. Evolution, 65: 3578–3589.

Ehrlich PR, Wilson EO. 1991. Biodiversity studies: Science and policy. Science, 253: 758–762.

Environmental Research Systems Institute (ESRI). 2010. World Continents from ESRI data and maps CD. Redlands, CA.

FitzJohn RG, Pennell MW, Zanne AE, Stevens PF, Tank DC, Cornwell WK. 2014. How much of the world is woody? Journal of Ecology, 102: 1266–1272.

Franklin J, Serra-Diaz JM, Syphard AD, Regan HM. 2017. Big data for forecasting the impacts of global change on plant communities. Global Ecology and Biogeography, 26: 6–17.

Gratton P, Marta S, Bocksberger G, Winter M, Trucchi E, Kühl H. in press. A world of sequences: can we use georeferenced nucleotide databases for a robust automated phylogeography? Journal of Biogeography.

Hansen MC, Potapov PV, Moore R, Hancher M, Turubanova S, Tyukavina A, Thau D, Stehman S, Goetz S, Loveland T et al. 2013. High-resolution global maps of 21st-century forest cover change. Science, 342: 850–853.

Hinchliff CE, Smith SA. 2014. Some limitations of public sequence data for phylogenetic inference (in plants). PLoS One, 9: e98986.

Hinchliff CE, Smith SA, Allman JF, Burleigh JG, Chaudhary R, Coghill LM, Crandall KA, Deng J, Drew BT, Gazis R et al. 2015. Synthesis of phylogeny and taxonomy into a comprehensive tree of life. Proceedings of the National Academy of Sciences, 112: 12764–12769.

Hortal J, de Bello F, Diniz-Filho JAF, Lewinsohn TM, Lobo JM, Ladle RJ. 2015. Seven shortfalls that beset large-scale knowledge of biodiversity. Annual Review of Ecology, Evolution, and Systematics, 46: 523–549.

James SA, Soltis PS, Belbin L, Chapman AD, Nelson G, Paul DL, Collins M. 2018. Herbarium data: Global biodiversity and societal botanical needs for novel research. Applications in Plant Sciences, 6: e1024.

Jones KE, Bielby J, Cardillo M, Fritz SA, O’Dell J, Orme CDL, Safi K, Sechrest W, Boakes EH, Carbone C et al. 2009. PanTHERIA: a species-level database of life history, ecology, and geography of extant and recently extinct mammals. Ecology, 90: 2648–2648.

Joppa LN, Roberts DL, Pimm SL. 2011. How many species of flowering plants are there? Proceedings of the Royal Society of London B: Biological Sciences, 278: 554–559.

Kattge J, Díaz S, Lavorel S, Prentice IC, Leadley P, Bönisch G, Garnier E, Westoby M, Reich PB, Wright IJ et al. 2011. Try - a global database of plant traits. Global Change Biology, 17: 2905–2935.

Keith DA. 2015. Assessing and managing risks to ecosystem biodiversity. Austral Ecology, 40: 337–346.

Li X, Yang Y, Henry RJ, Rossetto M, Wang Y, Chen S. 2015. Plant dna barcoding: from gene to genome. Biological Reviews, 90: 157–166.

Lomolino MV. 2004. Conservation biogeography. Frontiers of Biogeography: new directions in the geography of nature, pp. 293–296.

May RM. 2004. Tomorrow’s taxonomy: Collecting new species in the field will remain the rate-limiting step. Philosophical Transactions: Biological Sciences, 359: 733–734.

Mora C, Tittensor DP, Adl S, Simpson AG, Worm B. 2011. How many species are there on earth and in the ocean? PLoS Biol, 9: e1001127.

Nilsson RH, Wurzbacher C, Bahram M, Coimbra VRM, Larsson E, Tedersoo L, Eriksson J, Duarte C, Svantesson S, SÃąnchez-GarcÃηa M et al. 2016. Top 50 most wanted fungi. MycoKeys, 12: 29–40.

Pennell MW, FitzJohn RG, Cornwell WK. 2016. A simple approach for maximizing the overlap of phylogenetic and comparative data. Methods in Ecology and Evolution, 7: 751–758.

Pennell MW, FitzJohn RG, Cornwell WK, Harmon LJ. 2015. Model adequacy and the macroevolution of angiosperm functional traits. The American Naturalist, 186: E33–E50.

Pyke GH, Ehrlich PR. 2010. Biological collections and ecological/environmental research: a review, some observations and a look to the future. Biological Reviews, 85: 247–266.

RBG Kew. 2016. The State of the World’s Plants Report—2016. Royal Botanic Gardens, Kew.

Rodrigues AS, Pilgrim JD, Lamoreux JF, Hoffmann M, Brooks TM. 2006. The value of the IUCN red list for conservation. Trends in Ecology & Evolution, 21: 71–76.

Schrodt F, Kattge J, Shan H, Fazayeli F, Joswig J, Banerjee A, Reichstein M, Bönisch G, Díaz S, Dickie J et al. 2015. Bhpmf–a hierarchical bayesian approach to gap-filling and trait prediction for macroecology and functional biogeography. Global Ecology and Biogeography, 24: 1510–1521.

Sokal R, Rohlf F. 1981. The principles and practice of statistics in biological research. 2nd edn. Freeman, WH, New York.

Taugourdeau S, Villerd J, Plantureux S, Huguenin-Elie O, Amiaud B. 2014. Filling the gap in functional trait databases: use of ecological hypotheses to replace missing data. Ecology and Evolution, 4: 944–958.

Team eBird. July 2014. ebird’s missing species.

Theophrastus. 1916. Enquiry Into Plants, Translated by A.F. Hort. Harvard University Press.

Thiers B. 2017. The world’s herbaria 2016: A summary report based on data from index herbariorum. Website http://sweetgum.nybg.org/science/ih/ [accessed 9 September 2017].

Veiga AK, Saraiva AM, Chapman AD, Morris PJ, Gendreau C, Schigel D, Robertson TJ. 2017. A conceptual framework for quality assessment and management of biodiversity data. PloS one, 12: e0178731.

Wheeler QD, Raven PH, Wilson EO. 2004. Taxonomy: impediment or expedient? Science, 303: 285.

Wilman H, Belmaker J, Simpson J, de la Ros aC, Rivadeneira MM, Jetz W. 2014. Eltontraits 1.0: Species-level foraging attributes of the world’s birds and mammals. Ecology, 95: 2027–2027.

Wood S. 2017. Generalized Additive Models: An Introduction with R. 2nd edn. Chapman and Hall/CRC.

Younis S, Weiland C, Hoehndorf R, Dressler S, Hickler T, Seeger B, Schmidt M. 2018. Taxon and trait recognition from digitized herbarium specimens using deep convolutional neural networks. Botany Letters, 0: 1–7.

